# Graded, multi-dimensional intragroup and intergroup variations in primary progressive aphasia and post-stroke aphasia

**DOI:** 10.1101/2019.12.29.882068

**Authors:** Ruth U. Ingram, Ajay D. Halai, Gorana Pobric, Seyed Sajjadi, Karalyn Patterson, Matthew A. Lambon Ralph

**Affiliations:** Division of Neuroscience and Experimental Psychology, School of Biological Sciences, University of Manchester, UK; MRC Cognition & Brain Sciences Unit, University of Cambridge, UK; Department of Neurology, University of California, Irvine, Irvine, USA; Department of Clinical Neurosciences, University of Cambridge

**Author notes:** <u>Correspondence to</u>: Ruth U. Ingram, Division of Neuroscience and Experimental Psychology, School of Biological Sciences, Zochonis Building, University of Manchester, Brunswick Street, Manchester, M13 9PL, UK, or, Prof. Matthew A. Lambon Ralph, PhD.

**Keywords:** aphasia, stroke, neurodegeneration, classification

## Abstract

Language impairments caused by stroke (post-stroke aphasia) and neurodegeneration (primary progressive aphasia) have overlapping symptomatology, nomenclature and are classically divided into categorical subtypes. Surprisingly, primary progressive aphasia (PPA) and post-stroke aphasia (PSA) have rarely been directly compared in detail. Rather previous studies have compared certain subtypes (e.g., semantic variants) or have focussed on a specific cognitive/linguistic task (e.g., reading). This study assessed a large range of linguistic and cognitive tasks across the full spectra of PSA and PPA. We applied varimax-rotated principal component analysis to explore the underlying structure of the variance in the assessment scores. Similar phonological, semantic and fluency-related components were found for PSA and PPA. A combined principal component analysis across the two aetiologies revealed graded intragroup and intergroup variations on all four extracted components. Classification analysis was employed to test, formally, whether there were any categorical boundaries for any subtypes of PPA or PSA. Semantic dementia proved to form a true diagnostic category (i.e., within group homogeneity and distinct between group differences), whereas there was considerable overlap and graded variations within and between other subtypes of PPA and PSA. These results suggest that (a) a multi-dimensional rather than categorical classification system may be a better conceptualisation of aphasia from both causes, and (b) despite the very different types of pathology, these broad classes of aphasia have considerable features in common.

## Introduction

When stroke or neurodegeneration affects language-processing regions of the brain, patients can present with an acquired impairment of speech and language known as aphasia (McNeil and Pratt, 2001), or primary progressive aphasia (PPA) (Mesulam, 2001), respectively. To differentiate the two, we will refer to aphasia as a consequence of stroke as post-stroke aphasia (PSA). Two clinical and theoretical issues are addressed in this study. First, despite the similarity of symptoms and nomenclature in subtypes of PSA and PPA, there have been few – if any – detailed direct comparisons across the full range of PSA and PPA. Secondly, although diagnostic subtypes have been proposed for both forms of aphasia, patients often vary greatly within each category or commonly fall between classifications (and thus are referred to as ‘mixed’). This suggests that the phenotype differences observed across patients might reflect graded variations across multi-dimensional aphasic spectra rather than a series of mutually exclusive, coherent diagnostic categories (Lambon Ralph et al., 2003; Migliaccio et al., 2009; Ridgway et al., 2012; Stopford et al., 2008; Warren et al., 2012). By combining detailed assessment data across the full ranges of PSA and PPA, this study was able to map out these graded intergroup and intragroup variations.

Although arising from very different pathologies, PSA and PPA share symptomatology. Despite these clear superficial behavioural similarities, detailed direct comparisons between PSA and PPA are rare and thus it is still unclear, if the degree and nature of the symptoms are the same, or if the vocabulary terms used to describe the patients and their symptoms are truly equivalent. The small number of previous comparative studies have been focused on either specific tasks or linguistic/cognitive domains. For example, Patterson *et al.* (2006) compared speech production and phonological deficits in a selection of non-fluent subtypes of PSA and PPA. Jefferies and Lambon Ralph (2006) compared semantically-impaired PSA and PPA patients on a range of linguistic and non-linguistic semantic tasks (see also Jefferies et al., 2008). Thompson *et al.* (2013) compared syntactic processing in agrammatic and anomic forms of PSA and PPA (see also Budd et al., 2010; Faria et al., 2013; Thompson et al., 2012). Whilst these important studies have advanced our understanding of specific language features for selected subtypes of PPA/PSA, larger scale studies are needed for at least two reasons: (a) it is important to explore performance simultaneously across a broad spectrum of language and cognitive areas in order to situate and understand any one specific task; and (b) comparisons of select PPA/PSA subtypes make the assumption that the subtypes can be readily identified and are the most appropriate basis for the comparison.

People with PPA or PSA display considerable variation in the nature and severity of their impairments (e.g., naming, repetition, comprehension, reading, etc.) – but what is the basis of these variations? Behavioural variation in health or disease can be split into two types reflecting the presence of either multiple, mutually exclusive coherent categories of person/patient, or graded variations along different dimensions. All true categorical classification systems are based on two assumptions: 1) that there is homogeneity within each category or type; and 2) that there are distinct boundaries between categories (Schwartz, 1984). As is traditional in neurology and neuropsychology, categorical subtypes of PSA and PPA have been proposed. The Boston Diagnostic Aphasia Examination (Kaplan, 1983), for example, categorises PSA patients into one of six subtypes based on repetition, speech output fluency and comprehension (see also the Western Aphasia Battery (Kertesz, 2007)). Likewise, the consensus derived classification system for PPA (Gorno-Tempini et al., 2011) delineates three categorical subtypes: semantic dementia/semantic variant PPA, non-fluent/agrammatic variant PPA and logopenic variant PPA, though numerous additional subtypes are often proposed (such as agrammatic PPA without apraxia of speech (Tetzloff et al., 2019), or primary progressive apraxia of speech (Josephs et al., 2012)).

There is growing evidence, however, that a formal categorical approach is limited and does not capture the true nature of the patients’ variations. Thus, (a) rather than homogeneity within each category, there is significant variation (e.g., consider the different presentations of anomic aphasia or non-fluent progressive aphasia); (b) patients’ categorical membership can change (with recovery in PSA and decline in PPA); and (c) there are very fuzzy boundaries between categories (e.g., the boundary between conduction aphasia and Wernicke’s aphasia). One consequence is that in both PSA and PPA there is a considerable proportion of patients who must be classified as having ‘mixed’ aphasia because they either do not fulfil the criteria for any subtype, or even fulfil the criteria for more than one subtype (Benson, 1979; Botha et al., 2015; Gil-Navarro et al., 2013; Harris et al., 2013; Knibb et al., 2009; Matias-Guiu et al., 2014; Mesulam et al., 2008; Mesulam et al., 2012; Sajjadi et al., 2012a; Spinelli et al., 2017; Utianski et al., 2019; Wertz et al., 1984; Wicklund et al., 2014).

These limitations of categorical classification systems have long been understood clinically (Caramazza, 1984) (e.g., the limited use of broad ‘brush stroke’ classifications like Broca’s aphasia in describing an individual’s unique impairments for the purpose of therapy (Feyereisen et al., 1986; Gordon, 1998)) and have recently begun to be addressed from a theoretical perspective exploring an alternative, non-categorical way to conceptualise behavioural variations (Butler et al., 2014; Mirman et al., 2015a). These new approaches are based on the second source of individual differences noted above – namely, graded variations along continuous behavioural dimensions. Recent studies have reconceptualised the variations in PSA as forming an aphasic multi-dimensional space with each patient taking up a different position (typically varying in terms of phonology, semantics, speech fluency and, when assessed, non-language cognitive skills (Butler et al., 2014; Halai et al., 2017; Halai et al., 2018a; Schumacher et al., 2019). In this formulation the classical aphasia labels (e.g., conduction, Broca’s, etc.) do not represent categories per se, but rather are verbal pointers to a subregion in the multi-dimensional space. By way of analogy, one can think of patients as colour hues across the red, green and blue (RGB) colour space. It is possible to recognise clear differences (such as yellow (e.g., Broca) vs. blue (Wernicke), etc.) but also to capture the graded and unbounded variations between colours (e.g., there are many types of blue, its boundary with greens or violets is unclear, there are many hues (e.g., teal, maroon, etc.) that are hard to classify uniquely, and perceivers (cf. clinicians/researchers) have slightly different definitions for each colour (cf. clinical label)). Accordingly, some key aims of the current study were: (a) to test if the same approach can be applied to PPA (in contrast to other studies where methods capable of capturing graded variation have only been used as an intermediate step towards categorising proposed subtypes of PPA (Hoffman et al., 2017; Mesulam et al., 2009)); (b) to compare the multi-dimensional spaces for PSA and PPA; (c) to test if a single multi-dimensional space can be formed for PSA and PPA to allow direct, intragroup and intergroup comparisons. These aims were tackled through two large PSA and PPA cohorts (inclusive of typical and mixed cases), both completing large-scale, detailed neuropsychological and aphasiological test batteries.

## Methods

We initially applied principal component analysis (PCA) to PPA and PSA separately. This allowed us to compare qualitatively the resultant multi-dimensional space for each patient group without forcing the two groups into a single space. Given that the two group-specific PCA results were very similar in form, available PSA patients were re-assessed using a shared test battery derived from the PPA test battery, so that all patients could be entered simultaneously into a unified PCA. This enabled direct comparisons of both intergroup and intragroup variations.

### Patients

All patients were recruited non-selectively (with respect to subtype-level behavioural presentation) to sample the full space and severities of behavioural impairments in both PPA and PSA. Although diagnostic subtype labels were applied for descriptive purposes, the inability to apply a single diagnostic label was not grounds for exclusion in either cohort. Demographic details are shown in Table 1.

Seventy-six people with chronic PSA were prospectively recruited from community groups and speech and language therapy services in the North West of England. Patients were included if they reported a single left hemisphere stroke at least 12 months prior to assessment and were native English speakers. A portion of the PSA cases have been reported by Butler *et al.* (2014) and Halai *et al.* (2017) (31/70), and by Halai *et al.* (2018b) (70/76). All patients were classified into diagnostic subtypes by application of the Boston Diagnostic Aphasia Examination (Kaplan, 1983). All patients provided informed consent under approval from the North West Multi-Centre Research Ethics Committee, UK. Thirty-four of the 76 PSA cases were available for re-testing on the shared battery for the unified PCA on PSA and PPA.

Forty-six people with PPA were prospectively recruited from memory clinics at Addenbrooke’s Hospital, University of Cambridge (UK), as part of a longitudinal study of PPA (Sajjadi, 2013). These cases have previously been reported by Sajjadi *et al.* (2012a; 2014; 2012b; 2012c) and Hoffman *et al.* (2017). Patients with PPA were recruited on the basis of meeting the core criteria for PPA (Mesulam, 2001) then classified into a diagnostic subtype by application of the Gorno-Tempini *et al.* (2011) criteria or given the label ‘mixed PPA’ if unclassifiable. Exclusion criteria included other causes of aphasia (e.g., non-neurodegenerative pathology), non-native English speakers and any other neurological or major psychiatric illness. The PPA dataset comprised data from two longitudinal rounds of testing to assess change over time. On average, the second round of data was collected after 12.7 months (standard deviation: 0.9 months). Following Lambon Ralph *et al.* (2003), the participants who had scores for both rounds were treated as pseudo-independent observations. This resulted in a total of 82 data points for analysis. This approach was employed because PCA is a data-hungry method (Guadagnoli and Velicer, 1988) which benefits from having adequate sampling of as much of the potential PPA ‘space’ as possible. All patients, or next of kin where appropriate, provided informed consent under approval from the Cambridge Regional Ethics Committee.

### Neuropsychological assessments

#### Post-stroke aphasia test battery

The tests included in the PSA test battery are shown in Supplementary Figure 1, and described in Halai *et al.* (2017) and Butler *et al.* (2014). Briefly, the battery assessed connected speech, comprehension of grammar, auditory discrimination, repetition, semantic knowledge, naming, working memory and attention/executive function.

#### Primary progressive aphasia test battery

The tests included in the PPA test battery are shown in Supplementary Figure 2, and described by Sajjadi et al (2013). Briefly, this battery assessed connected speech, comprehension of grammar, grammatical ability in sentence production, repetition, semantic knowledge, naming, phonological discrimination, working memory, attention and executive function, visuospatial skills, and oro-buccal and limb praxis.

#### Shared battery

In order to establish the shared multi-dimensional space of PSA and PPA, available PSA cases were re-tested on a shared test battery which was derived from the PPA test battery. Thirty-three tests were including in the shared battery, shown in Supplementary Figure 3. This battery assessed attention and executive function, repetition, sentence comprehension and production, semantic memory, visuospatial skills, praxis, connected speech, naming, and phonological discrimination.

#### Data analysis

All raw behavioural scores were converted to percentages. For measures without a fixed maximum score, scores were converted to a percentage of the maximum score across the relevant cohort or both cohorts for the unified PCA. Missing data were imputed using probabilistic principal component analysis (PPCA) (Ilin and Raiko, 2010). This approach was chosen as the results were stable when compared to versions of the analyses without imputation (i.e., list-wise exclusion analysis). PPCA requires that the number of components to be extracted is specified *a priori,* so a k-fold cross validation approach (Ballabio, 2015) was used to choose the number of components giving the lowest root mean squared error for held-out cases over 1000 permutations. This approach was also used to select the optimal number of components for subsequent PCA using the imputed dataset. The imputed datasets were entered into PCAs (conducted in SPSS 23), with varimax rotation to aid cognitive interpretation of the extracted dimensions. The adequacy of the sample size for each PCA was determined using Kaiser Meyer Olkin measure of sampling adequacy.

The separate PCAs for PPA and PSA could not be compared directly since they did not share the same tasks, so they were compared qualitatively by analysing the type of tasks that loaded most heavily onto each extracted component. This approach was also used to compare the separate PCAs to the unified PCA. However, since the unified PCA was conducted on data from both groups on the shared battery, this made direct, quantitative, intra- and inter-group comparisons possible. To put the relative regions of the multi-dimensional space occupied by PSA and PPA into perspective, control norms were projected into the unified PCA space by normalising to the patient mean and standard deviation, and then using the factor coefficients to generate factor scores for an average control participant.

Finally, formal analyses were conducted to test for the presence of subgroup categories (i.e., subgroups with relatively high intra-group homogeneity and distinct inter-group differences). If one or more subgroups formed a true category, they would be represented in the PCA multi-dimensional space as a homogenous group of data points, and it would be possible to define formal diagnostic boundaries with the other subgroups in terms of cut-off scores on each extracted dimension. To investigate this, formally, the unified PCA was systematically swept to find the combination of cutoff values across all dimensions that gave the highest sensitivity index (d prime – d’) value per diagnostic subtype within aetiology. PCA solutions are always scaled using z-scoring, and in this study the dimensions ranged from approximately −3 to +2 with zero representing the centroid of the patient cohort. These dimensions were swept iteratively at intervals of 0.05. The d’ equation was adapted to account for extreme values (0 or 1) for the rate of hits or false alarms (Macmillan and Kaplan, 1985), resulting in the maximum d’ value achievable being 4.65; thus a d’ value near 4.65 would be suggestive of distinct categorical boundaries and within-group homogeneity. To establish the likelihood of achieving these d’ values by chance, diagnostic group-membership was randomised within aetiology and d’ re-calculated over 10,000 iterations to give a distribution of d’ values.

The combination of cut-off values along each dimension that gave the maximum d’ value (i.e., highest possible sensitivity) were then treated as diagnostic ‘criteria’ for new data-driven diagnostic groups. The hits from the d’ analysis represent cases whose factor scores correctly met the cut-off values for their own data-driven diagnostic group. The false alarms represent cases whose factor scores incorrectly met the cut-off values for any other data-driven diagnostic group, i.e., cases who were misclassified. These misclassifications occurred despite the cut-offs representing the best possible (highest sensitivity) between-group boundaries that could be found in the iterative sweep through the entire multi-dimensional space. The distribution of misclassifications amongst subtypes of each aetiology was extracted from the false alarms associated with each data-driven diagnostic group. It is important to note that it was possible for a single case’s factor scores to meet the cut-off values for more than one data-driven diagnostic group simultaneously (or none), and this may or may not have included their own group.

## Results

### Demographics

**Table 1-.** demographic details per subtype of the post-stroke aphasia and primary progressive aphasia cohorts (data presented as Mean (standard deviation).

| Group | Subtype | N (F) | Age | Education (years) | Time with aphasia (years) |
| --- | --- | --- | --- | --- | --- |
| PSA | Anomia | 30 (11) | 63.8 (13.4) | 12.4 (2.7) | 4.2 (4.3) |
|  | Broca | 13 (1) | 62.7 (13.0) | 11.9 (1.8) | 4.3 (3.4) |
|  | Conduction | 4 (1) | 62.0 (10.7) | 13.8 (3.2) | 1.8 (0.9) |
|  | Global | 9 (0) | 66.4 (9.0) | 11.3 (0.7) | 5.9 (4.6) |
|  | Mixed non-fluent | 16 (4) | 68.1 (9.0) | 11.5 (1.0) | 6.3 (4.9) |
|  | TMA | 2 (1) | 74.5 (2.1) | 11.0 (0.0) | 6.8 (4.1) |
|  | TSA | 1 (0) | 63.0 | 12.0 | 2.0 |
|  | Wernicke/conduction | 1 (1) | 77.0 | 16.0 | 2.8 |
| PPA | Logopenic | 2 (1) | 71.0 (4.2) | 11.0 (2.8) | 2.0 (0.0) |
|  | Mixed PPA | 16 (12) | 72.7 (5.2) | 10.8 (1.9) | 3.3 (1.4) |
|  | PNFA | 12 (7) | 69.3 (7.3) | 13.0 (3.8) | 3.2 (1.4) |
|  | SD | 16 (8) | 67.1 (8.5) | 13.9 (3.3) | 4.2 (1.3) |

### Principal components analysis

#### Post-stroke aphasia

The PCA for the PSA cohort was robust (Kaiser Meyer-Olkin = 0.84) and produced a 4-factor rotated solution which accounted for 76.7% of variance in PSA patients’ performance (F1 = 30.4%, F2 = 17.5%, F3 = 15.2%, F4 = 13.7%). The factor loadings of each behavioural assessment onto the extracted components are shown in Supplementary Figure 1.

Measures loading heavily onto the first factor were tests of repetition (PALPA words, non-words), naming (Boston, Cambridge), phonological working memory (digit span), auditory comprehension (CAT sentence comprehension), and phonological sensitivity (PALPA minimal pairs). These tests all require phonological processing; hence we called this factor ‘Phonology’.

Measures that loaded heavily onto the second factor were tests of discrimination (PALPA minimal pairs), matching (word-to-picture matching), fluency and rule detection (Brixton, Raven’s). Whilst spanning different aspects of language and cognition, these tests all share the feature of having high executive demands; hence, we called this factor ‘Executive Function’.

Measures with strong loadings onto the third factor included speech rate (words per minute) and speech quanta (total number of words). We called this factor ‘Speech Production’.

Measures with strong loadings onto the fourth factor were tests of semantic knowledge (synonym judgment, word-to-picture matching) and semantic richness of speech (mean length per utterance). Furthermore, measures of naming (Boston, Cambridge) and sentence comprehension (CAT) also had moderate factor loadings onto this factor. These tests all require semantic knowledge; hence we called this factor ‘Semantics’.

#### Primary progressive aphasia

The PCA for the PPA cohort also generated a robust result (Kaiser Meyer-Olkin=0.85) with a 5-factor rotated solution which accounted for 72.4% of variance (F1 = 23.7%, F2 = 17.8%, F3 = 14.5%, F4 = 9.8%, F5 = 6.7%). The factor loadings of each behavioural assessment onto the extracted components are shown in Supplementary Figure 2.

The tests loading onto the first factor all required retaining phonological information (e.g. single digits, words, numbers, whole sentences) in mind; hence we called this factor ‘Phonological Working Memory’. These measures included tests of phonological sensitivity (non-word minimal pairs), attention and executive function (digit span forwards and backwards, letter span similar and dissimilar phonemes), repetition (words, non-words and sentences), sentence comprehension (SECT, TROG) and cube counting (VOSP).

The second factor comprised heavy loadings from tests relying on semantic knowledge; hence we called this factor ‘Semantics’. These tasks included tests of semantic knowledge (Cambridge naming, Point from Repeat and Point, Category Fluency), semantic association (Camel & Cactus), recognition of irregular words, and sentence comprehension (SECTV).

The third factor was characterised by strong loadings from measures of speech rate (words per minute) and speech quanta (total number of words), in addition to oro-buccal praxis. A test of executive function requiring drawing and counting (TMT-A) also had high loadings (note, patients often count under their breath or out loud to complete the TMT-A). Accordingly, this factor appeared to capture the motor aspect of speech, hence we called this factor ‘Motor Speech Production’.

Visuospatial tests of executive function loaded heavily onto the fourth factor. Specifically, tests of switching (TMTB), counting and visual imagery (VOSP), and copying and visuospatial recall (Rey Complex Figure) had high loadings on this factor, hence we called this factor ‘Visuo-Executive Function’.

Loadings onto the fifth factor were dominated by tests of sentence production (MAST), measures of semantic richness of speech (mean length per utterance) and generation of items (letter fluency from the Addenbrooke’s Cognitive Examination – Revised (Mioshi et al., 2006)). Having accounted for motor speech production, semantics and executive demands in earlier factors, the remaining aspect of these tests which might be captured in this final independent factor could be the generative aspect of speech production. Hence, we called this factor ‘Speech Generation’.

#### Unified PCA on the shared battery

Given that the two group-specific batteries and PCAs generated similar types of dimensions (phonology, semantics, executive skill and aspects of speech production), a formal direct comparison through a shared battery and single PCA spanning both groups was both merited (i.e., there was *prima facia* evidence of shared symptoms and variations) and permitted formal intra-group and inter-group comparisons by enabling inclusion of all patients across a single multi-dimensional space. The unified PCA was again robust (Kaiser Meyer-Olkin=0.88), with a 4-factor rotated solution accounting for 67.4% of variance in patients’ performance (F1 = 23.5%, F2 = 16.6%, F3 = 14.8%, F4 = 12.6%), and bore a strong relationship with the factors identified in the group-specific test batteries. The factor loadings of each behavioural assessment onto the extracted components are shown in Supplementary Figure 3.

The first factor had high loadings from tests of repetition (words, non-words, sentences), phonological sensitivity and attention (digit and letter spans, non-word minimal pairs), and sentence comprehension (auditory and visual). These tests all require intact phonological processing, hence we called this factor ‘Phonology’.

There were high loadings on the second factor for tests of semantic knowledge (Cambridge naming, pointing from the Repeat and Point test), generation of items in a semantic category (category fluency from the ACE), sentence comprehension (SECT) and address recall and recognition from the ACE. Hence, we called this factor ‘Semantics’.

Tests of attention and executive function in the visuospatial domain (VOSP, TMT, Rey Complex Figure) all loaded heavily onto the third factor. As above, we called this factor ‘Visuo-Executive Function’.

The fourth factor had high loadings from measures of speech quantity (words per minute, total number of words and mean length per utterance). Measures of praxis (Oro-buccal and limb) and phonological working memory (digit span backwards) also had high loadings onto this factor. Given that phonological ability and executive functions have been accounted for already, this factor probably captured the speech production element of the digit span test (patients often repeat the string of digits to themselves before reporting them backwards). These tests therefore all require production of speech, and coupled with the loadings from the praxis tests, we interpreted this as a ‘Motor Speech Production’ factor.

#### Inter-group comparisons

To illustrate the components extracted in each PCA, exemplar tests with strong and relatively unique loadings onto each factor across the three PCAs are plotted together in Figure 1 (the full plots of all tests loadings on all PCA factors are shown in the Supplementary Figures). The specific example test chosen differed across PCAs due to the different test batteries, but where possible the same or a similar measure was chosen. This Figure highlights the similarity of the components extracted for both forms of aphasia, whether separately or in combination.

**Figure 1-.**
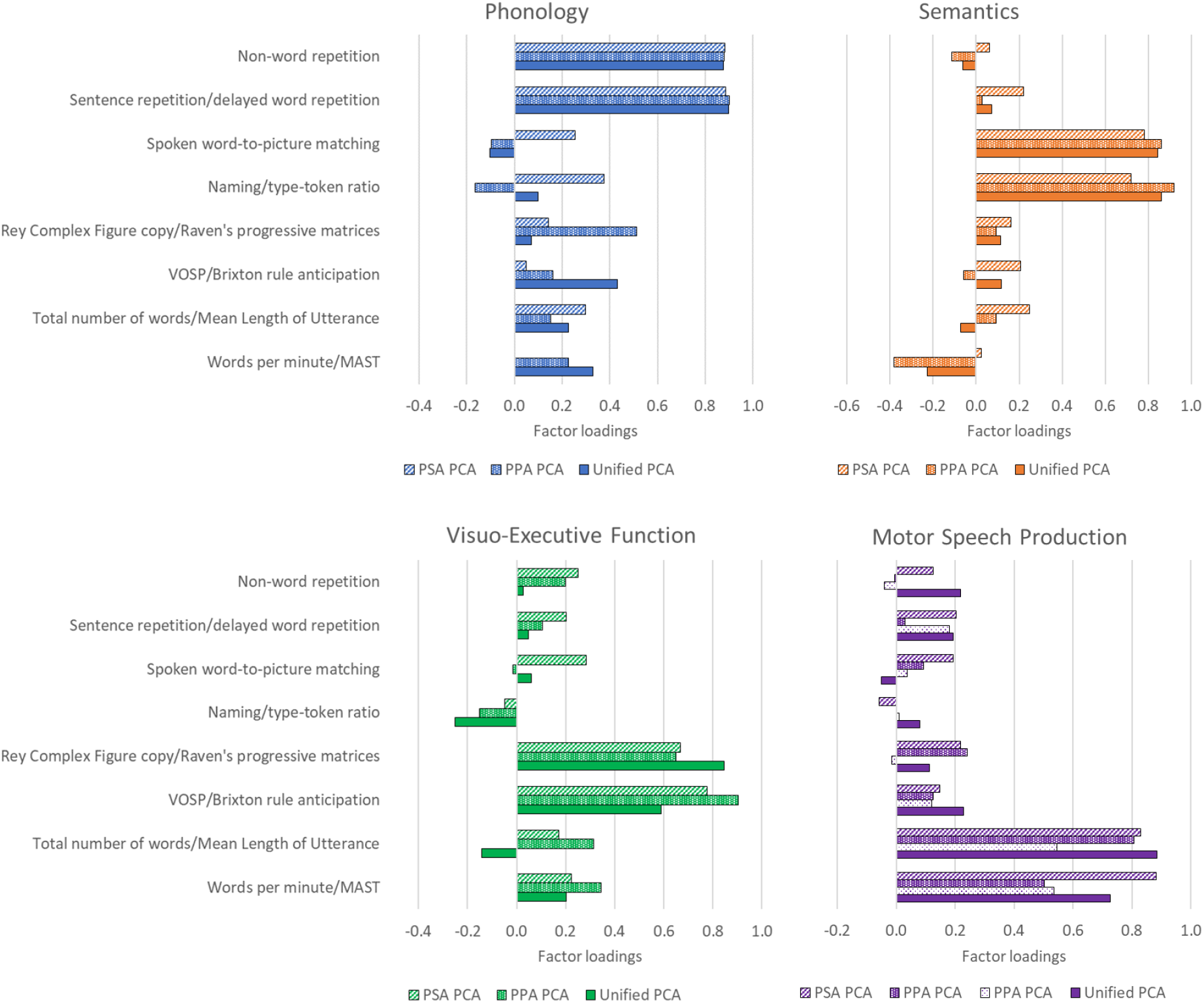
Inter-group comparison of the underlying dimensions of variance in PSA and PPA. Bars represent the factor loadings of exemplar tests onto each extracted factor. Factor loadings represent the weighting of each test on each factor and were used to suggest cognitive interpretations of the factors. The patterns of the bars represent the different PCAs; the PPA PCA extracted two speech production components, which are shown in different patterns on the Motor Speech Production panel. Abbreviations: PSA – post-stroke aphasia; PPA – primary progressive aphasia; PCA – principal component analysis; VOSP – Visual Object and Space Perception battery (Warrington and James, 1991), MAST – Make a Sentence Test (Billette et al., 2015).

Direct inter- and intra-group comparisons were possible in the shared multi-dimensional space of the unified PCA. Figure 2 plots the patients and their aphasia classifications (PSA in blue and PPA in red markers) into the four-dimensional factor space (Panel A maps the phonology and semantics factors, Panel B speech production vs. visuo-executive skill factors). Four key observations can be gleaned from these scatterplots: (i) intra-group graded differences: for both PPA (except semantic dementia, see below) and PSA there is considerable variation across cases within each subtype of aphasia and also overlap between the groups (e.g., conduction and anomic aphasia or PNFA and mixed PPA); (ii) inter-group differences: with regards to semantics and phonology the PSA and PPA cases are fully overlapping reflecting the clinical observation that the two aetiologies share many language symptoms; (iii) the two aetiologies are strongly separated in terms of speech fluency and co-occurring visuo-executive skills with the PSA cases dominating the space denoting poorer fluency yet better visuo-executive skills (lower right quadrant in Panel B). All forms of PSA (even those referred to as “fluent”) were less fluent that the PPA patients (with the exception of the most severe PNFA and mixed cases), whilst only the SD subset were able to match the PSA on visuo-executive skills; (iv) by eye, the only group which might form a coherent and separated cluster (cf. a true category) are those with SD (yellow crosses) in that they appear to uniquely occupy the combination of moderate-to-severe semantic impairment with good phonology (i.e., lower right quadrant in Panel A) and good visuo-executive function and speech fluency (i.e., upper right quadrant in panel B). We tested formally whether SD and any other groups form a true category in the subsequent analysis.

**Figure 2-.**
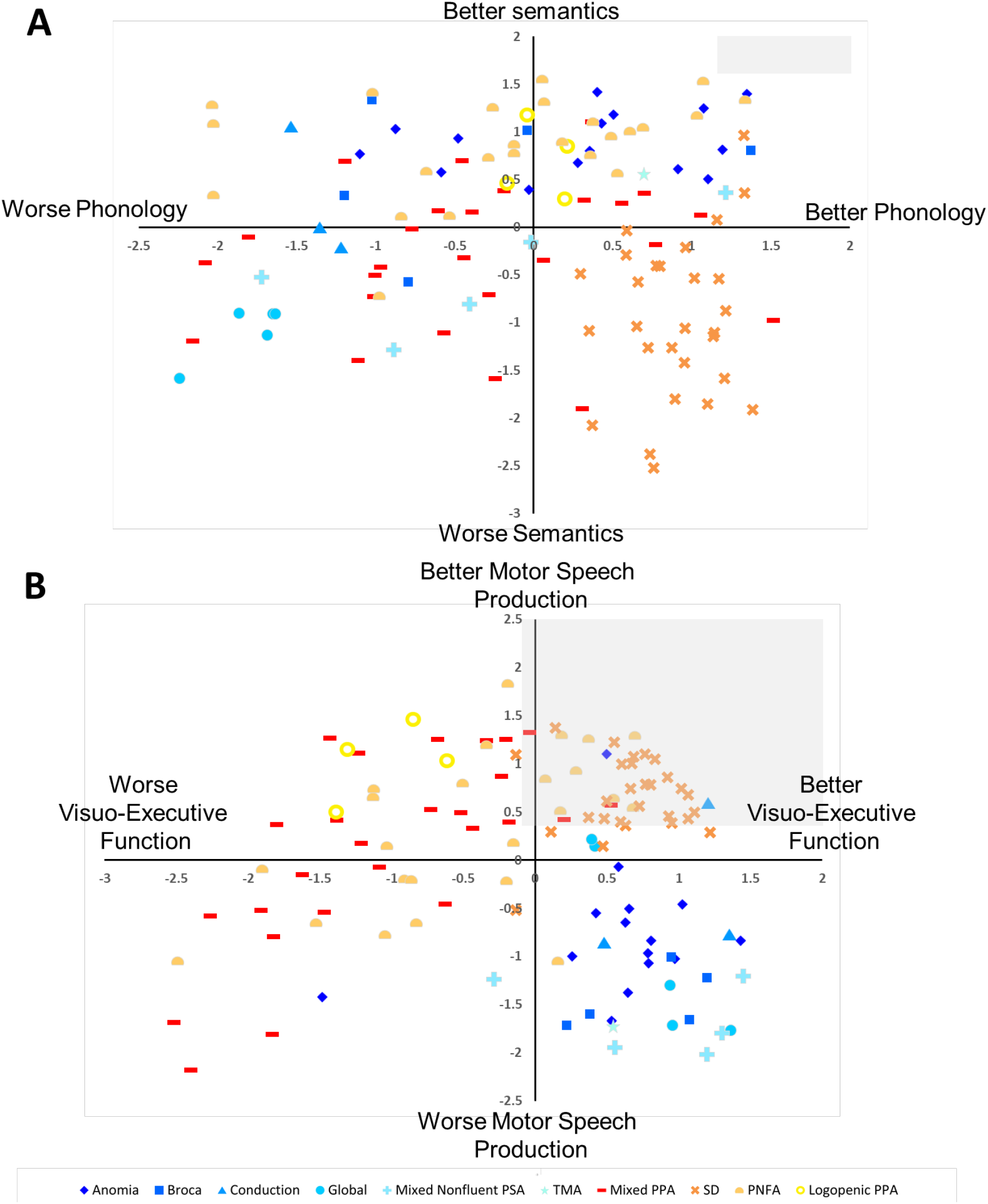
Regions of the shared multi-dimensional space of PSA and PPA occupied by each diagnostic subtype. Factor scores of all patients were plotted along all pairs of components extracted from the unified PCA. The origin is the mean of all patients. The factor scores are an expression of how many standard deviations a patient’s performance is from the group mean. The region of space reflecting preserved performance was calculated by projecting control norms into the patient space and is shaded in grey. PSA subtypes are blue-spectrum colours, PPA are red-spectrum colours. Abbreviations: PSA – post-stroke aphasia; PPA – primary progressive aphasia; TMA – transcortical motor aphasia; SD – semantic dementia; PNFA – progressive non-fluent aphasia.

#### Intra-group graded variation

For each subtype within each aetiology, the best combination of ‘diagnostic’ values across all four dimensions was derived using a data-driven search (see Methods; all values are displayed in Supplementary Table 1). These values were treated as cut-offs defining new data-driven diagnostic groups, which were labelled according to the subtype from which the cut-offs were derived. An illustration of the data-driven diagnostic cut-offs for SD is shown in Figure 3.

**Figure 3-.**
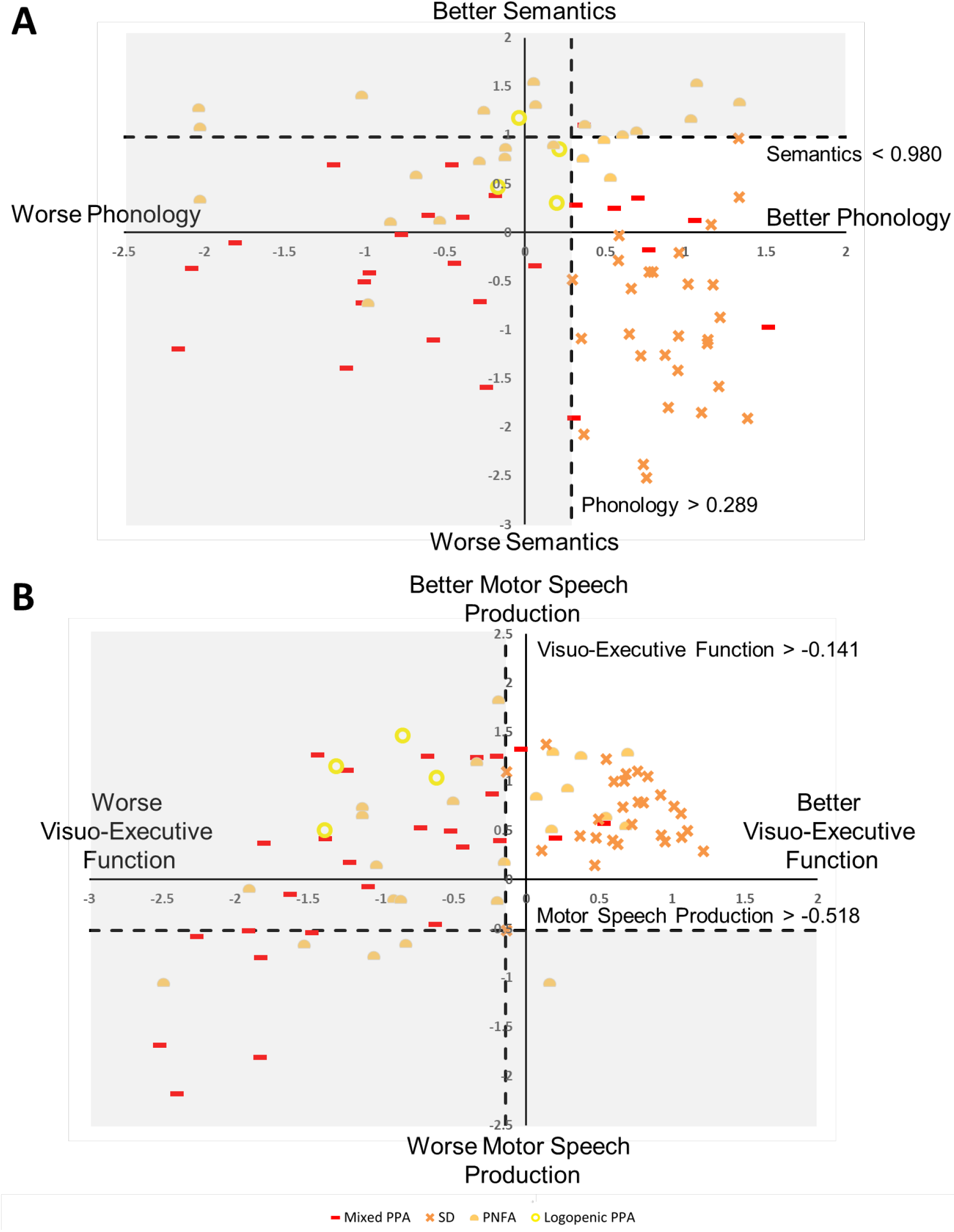
Data-driven diagnostic cut-off values for semantic dementia. Using a data-driven sweep at intervals of 0.05 through the entire four-dimensional space, the combination of cut-off values giving optimum sensitivity for semantic dementia was isolated. Applying the simultaneous combination of these four-dimensional cut-off values as diagnostic criteria gave perfect selectivity for semantic dementia. This implies that semantic dementia shows within-group homogeneity and distinct between-group differences, suggestive of a true diagnostic category. This process was repeated for all subtypes of PPA and PSA within each aetiology (cut-off values and d’ values per subtype in Supplementary Table 1). Abbreviations: SD – semantic dementia; PNFA – progressive non-fluent aphasia.

The pattern of hits and misclassifications associated with each combination of diagnostic cut-offs for PPA subtypes is shown in Table 2. The pattern of hits and misclassifications for PSA subtypes is shown in Supplementary Table 2, as these results will need validating in a larger cohort; in order to include a heterogeneous cohort reflecting the true phenotypic space of PSA, subtypes of PSA were included in this study even if they comprised only a single case. However, in terms of assessing whether the subtypes of PSA meet the assumptions of a true category, larger sample sizes will be needed to fully answer this question.

**Table 2-.** Distribution of misclassifications between clinical and data-driven diagnostic PPA groups. The cut-off values giving optimum sensitivity for each diagnostic group were treated as data-driven diagnostic criteria. Rows represent ‘real’ clinical diagnostic categories. The ‘Hits’ column represents the percentage of patients meeting the data-driven cut-off values for their own data-driven diagnostic group. The columns under ‘Misclassifications’ represent the percentage of cases whose factor scores (a) met the cut-off values for a different data-driven diagnostic group; (b) did not meet the cut-off values for any of the data-driven diagnostic groups; (c) met the cut-off values for more than one data-driven diagnostic group. These ‘Misclassifications’ columns are not mutually exclusive, so row totals do not add up to 100%.

|  |  | Data-driven diagnostic groups |  |  |  |  |  |  |
| --- | --- | --- | --- | --- | --- | --- | --- | --- |
|  |  | Hits | Misclassifications |  |  |  |  |  |
|  |  |  | lvPPA | mPPA | PNFA | SD | None | >1 |
| Clinical diagnostic groups (N) | lvPPA (4) | 100.0 | / | 50.0 | 100.0 | 0.0 | 0.0 | 100.0 |
|  | mPPA (26) | 92.3 | 0.0 | / | 38.5 | 0.0 | 3.8 | 34.6 |
|  | PNFA (24) | 91.7 | 4.2 | 20.8 | / | 0.0 | 4.2 | 16.7 |
|  | SD (28) | 100.0 | 0.0 | 25.0 | 3.6 | / | 0.0 | 28.6 |

The data presented in Table 2 are the percentages of patients, from each clinical diagnostic subgroup of PPA, whose factor scores met the data-derived cut-off values for each diagnostic group. The rows represent the ‘real’ clinical diagnostic categories for patients in this study. The ‘Hits’ column represents the percentage of patients meeting the data-driven cut-off values for their own diagnostic group. The columns under ‘Misclassifications’ represent the percentage of cases whose factor scores (a) met the cut-off values for a different (i.e., incorrect) diagnostic group; (b) did not meet the cut-off values for any of the possible data-driven diagnostic groups; (c) met the cut-off values for more than one data-driven diagnostic group (e.g., their own group and one other group). These ‘Misclassifications’ columns are not mutually exclusive and thus cases falling into more than one classification are tabulated in the ‘>1’ column; consequently, the row totals do not add up to 100%.

For example, the optimum cut-off values for SD were highly selective for SD, with 100% of the SD cases’ factor scores meeting these values (‘Hits’ column). Furthermore, as can been seen from the ‘Misclassifications – SD’ column, there were no misclassifications of patients from other diagnostic groups as SD. This corresponds to the highest d’ value of 4.46 (p < .001) for SD and suggests that the SD cases from which the data-driven diagnostic criteria were derived show within-group homogeneity and clearly distinct between-group boundaries. This corroborates our earlier qualitative interpretation of SD as occupying a unique area in the multi-dimensional space from the Unified PCA. Due to the data-driven criteria for other PPA subtypes being less selective for their target subtype, some SD cases were misclassified (columns ‘Misclassifications – mPPA’ and ‘Misclassifications – PNFA’); accordingly, in the ‘Misclassifications – >1’ column, 28.6% of the SD cases met both the SD cut-offs and the cut-offs for either mPPA or PNFA.

The lvPPA data-driven diagnostic criteria (d’ = 3.38, p < .001) had a perfect hit rate but some misclassifications of PNFA cases, resulting in a lower d’. As with the subtypes of PSA with small sample sizes, the results for lvPPA will benefit from replications in future studies with larger samples. The data-driven diagnostic criteria for PNFA (d’ = 2.03, p < .001) were much less selective as not only did they incur misclassifications of all other subtypes, but they also failed to classify all the PNFA cases correctly (hit rates around 90%). This shows that even the optimal data-driven diagnostic criteria for PNFA were insufficiently selective, implying that PNFA cases do not display within-group homogeneity and distinct between group boundaries like SD.

By definition, the mPPA group represent cases who did not meet the criteria for a single proposed subtype of PPA. Consistent with this clinical categorisation, the data-driven diagnostic criteria for this group (d’ = 2.10, p <.001) also showed low selectivity. The mPPA criteria failed to capture all the mPPA cases and incorrectly captured cases from all other subtypes.

## Discussion

This study had two principal aims: (i) to undertake a large-scale direct comparison of PPA and PSA utilising detailed aphasiological and neuropsychological test batteries, and (ii) to reconsider the phenotype differences across patients with PSA and PPA in terms of graded variations along multiple principal dimensions.

The results confirm that there is meaningful, coherent structure in the language-cognitive variations across PPA and PSA patients. Rather than conceptualising such variations as mutually-exclusive categories, the results indicate that the patients’ variations reflect multiple, continuous, graded dimensions. This alternative approach has multiple advantages: (a) it is able to capture the patterns of overlap between different ‘subtypes’ of PPA and PSA (e.g., overlap in phonological impairments in many PPA and PSA cases) as well as their clear differences; (b) it captures the variations in performance within each ‘type’ of PPA and PSA; and (c) it meaningfully situates the ‘mixed’ aphasic patients alongside the other cases to generate a complete clinical picture of PPA and PSA. This is important given that there are high numbers of ‘mixed’ cases in everyday clinical practice. It is, perhaps, important to note that these dimensions are not new categories but rather each patient represents a specific point in the graded multidimensional space.

The cognitive and language impairments in PSA and PPA, both when considered in isolation and when considered in a single unified framework, could be captured by four main dimensions of underlying variation: phonology, semantics, speech production/motor output fluency, and executive-cognitive skill. This finding is a direct replication of what has been found previously for this PSA cohort (Butler et al., 2014; Halai et al., 2017) and by numerous international groups (Kümmerer et al., 2013; Lacey et al., 2017; Mirman et al., 2015a; Mirman et al., 2015b; Tochadse et al., 2018), and found to be statistically stable across different sample sizes and assessment batteries (Halai et al., 2018a). These studies have used lesion-symptom mapping methods to show that the principal components are associated with neural correlates that support the labels applied (e.g. components labelled ‘Phonology’ having neural correlates in left posterior perisylvian regions such as superior temporal gyrus, which have previously been shown to be involved in phonological processing). Future studies will be able to explore how the PCA dimensions observed in PPA map to the gradedly-varying patterns of atrophy across brain regions.

The fact that the same underlying dimensions were found for PPA as well as PSA indicates that these dimensions might reflect core “primary systems” for language activities (Patterson and Lambon Ralph, 1999; Ueno et al., 2014; Woollams et al., 2018). Past work has associated these primary systems with different brain areas: phonological processing and working memory with posterior superior temporal lobe and supra-marginal gyrus (Paulesu et al., 1993); semantic representation with anterior temporal lobe (ATL) (Lambon Ralph et al., 2017; Patterson et al., 2007); speech programming and fluency with premotor cortex and key underpinning white matter pathways (Basilakos et al., 2014); and executive functions with frontoparietal networks (Jurado and Rosselli, 2007; Marek and Dosenbach, 2018). As these regions can be affected in both middle cerebral artery PSA (Phan et al., 2005) and PPA (Gorno-Tempini et al., 2004), the similarity of their phenotypic spectra could reflect varying degrees of impairment to these core primary systems.

Plotting all patients’ factor scores into the shared multi-dimensional space showed that the non-SD PPA and PSA cases occupied an almost completely overlapping region of the Phonology-Semantics space. This contrasts with the SD cases who occupied an exclusive region of the multi-dimensional space, signifying their selective semantic impairment in the context of relatively preserved phonological abilities, coupled with motor speech production and executive function that are comparable to healthy controls. This might reflect the fact that SD arises from atrophy in extra-sylvian, ATL regions (Hodges et al., 1992; Rosen et al., 2002; Snowden et al., 1989), whereas the other forms of PPA and PSA are associated with damage to perisylvian regions (Grossman and Irwin, 2018).

In the space corresponding to Visuo-Executive Function vs. Motor Speech Production, there was separation of the two aetiologies. PSA patients occupied the region signifying less fluent speech production combined with relatively unimpaired visuo-executive ability. Most PPA patients showed the reverse pattern, although some PNFA and mPPA cases showed the combination of poor fluency and poor executive function. This separation is clinically interesting and important as it indicates that certain symptom terms – e.g., fluency – are not used in the same way across patient types; thus, many non-fluent progressive aphasics were more fluent than the “fluent” PSA cases (e.g., anomia and conduction aphasics). This result supports the findings of Patterson *et al.* (2006) who found that non-fluency in PSA and in PPA were not equivalent. Furthermore, the separation in terms of Visuo-Executive Function could reflect an aetiology-driven difference in damage to the multi-demand frontoparietal executive system (Marek and Dosenbach, 2018), and posterior cingulate and other medial regions. These regions support executive function and attention (Jurado and Rosselli, 2007) and are situated at the edges/outside of the MCA-perfused regions (Phan et al., 2005).

In addition to facilitating inter-group comparisons, the PCA method revealed graded intra-group differences. The subtypes of non-SD PPA and PSA occupied only partially differentiated positions within the four-dimensional space, with considerable variation within each “subtype” and overlap of cases across subtypes. This could reflect the overlapping atrophy/lesions in and around the perisylvian language regions in these forms of aphasia. Again, these findings indicate that phenotypic variations in non-SD PPA and PSA are unlikely to reflect different categories but rather graded variations along these dimensions. These graded differences can only be accounted for in categorical classification systems by using ‘mixed’ classifications (Sajjadi et al., 2012a; Wertz et al., 1984; Wicklund et al., 2014), but the methods in the current study were able to account for graded variation in a single multi-dimensional framework comprising four, clinically-intuitive underlying dimensions.

Based on this framework, the current diagnostic subtype labels can be reconceptualised as pointers towards particular regions of the multi-dimensional space, rather than labels for mutually-exclusive clinical categories. This approach does not preclude the fact that some labels might be pointers for more exclusive regions of space (e.g., SD or global PSA) than others (e.g., anomia or PNFA).

In fact, the concept of ‘SD’ seems to be a uniquely useful pointer for the exclusive region of the multi-dimensional space occupied by these cases. This aligns with (i) the original descriptions of SD, in particular the selective nature of their semantic impairment (Hodges et al., 1992; Snowden et al., 1989; Warrington, 1975), and (ii) previous work showing that SD is distinct from other forms of PPA; Bisenius *et al.* (2017) found that SD was the most readily differentiable subtype of PPA using Support Vector Machine approaches to evaluate the consensus criteria for PPA. Hoffman *et al.* (2017) applied k-means clustering to behavioural data in PPA and found that of their three-cluster solution, only one cluster was selective for a particular subtype of PPA and this was the SD cluster. Furthermore, by plotting PSA and PPA in the same space we provide support for previous work showing that semantic impairments in SD are unlike those found in PSA (Jefferies and Lambon Ralph, 2006; Lambon Ralph et al., 2017). This result probably reflects the fact that the distribution of damage in SD is distinctly different from those in non-SD PPA and PSA phenotypes. SD cases have hypometabolism and atrophy centred on the ATL bilaterally (Mummery et al., 2000), which data from other methods in healthy participants and patient groups has shown to be a key region for the formation of coherent concepts (Lambon Ralph et al., 2010). This finding agrees with Sajjadi *et al.* (2017) who found that the atrophy patterns for SD were more easily distinguishable (high sensitivity and specificity) than the other forms of PPA.

Describing the symptomatology of PSA and PPA in terms of gradedly-different regions within multi-dimensional space has a number of potential clinical implications. First, this approach allows us to begin to determine both the range and type of variations that are associated with each of the preexisting clinical labels, rather than reserving the use of each diagnostic label to a single, invariant prototypical pattern. Secondly, by extension, it also allows us to establish when and why certain subtypes of PPA/PSA are most likely to be confused with each other. Thirdly, it provides a single unifying framework within which both established and “mixed” aphasias can be considered. Fourth, future clinical research can explore whether considering the phenotype variations along continuous, dimensions (i.e., a transdiagnostic approach) rather than porous categorical systems might reveal clearer relationships between phenotype and atrophy, pathology or genetic markers. For example, past work in PSA has shown that utilising raw individual test scores or PSA categories leads to undifferentiated lesion correlates reflecting the whole MCA territory rather than specific subregions. When the same lesion-mapping analyses are repeated using the PCA-derived dimensions then much more discrete and interpretable subregions are revealed (cf. Butler et al., 2014). These clearer symptom-lesion maps can then be inverted in order to generate lesion-based diagnostics and prediction models (Halai et al., 2018). Finally, taking this multi-dimensional approach could inform a transdiagnostic selection process for treatment, therapy or clinical trials; in order to select a group of patients with relatively homogeneous behavioural symptoms, one could select patients who occupy a shared region of the multi-dimensional space (thereby sharing symptomatology across the core language systems captured by the dimensions), irrespective of their clinical diagnosis. Indeed, the importance of a transdiagnostic approach has been highlighted for Frontotemporal Lobar Degeneration (FTLD) with regards to shared apathy and impulsivity symptomatology (Lansdall et al., 2017; Lansdall et al., 2019; Passamonti et al., 2018).

In conclusion, we have shown that the internal structure of variation in PSA and PPA, in isolation or in a unified framework, can be captured with the same four underlying language-cognitive dimensions. Furthermore, semantic dementia appears to represent a robust diagnostic category, whilst patients with other forms of PPA and PSA might be better described in terms of their gradedly-different positions along these four principal language-cognitive dimensions.

## Acknowledgements

We thank all of the patients, their families and carers for their continued support of our research programmes. This research was supported by The Rosetrees Trust (A1699) and ERC (GAP: 670428 - BRAIN2MIND_NEUROCOMP).

