## Supplementary material for "Graded, multi-dimensional intragroup and intergroup variations in primary progressive aphasia and post-stroke aphasia"

### Supplement

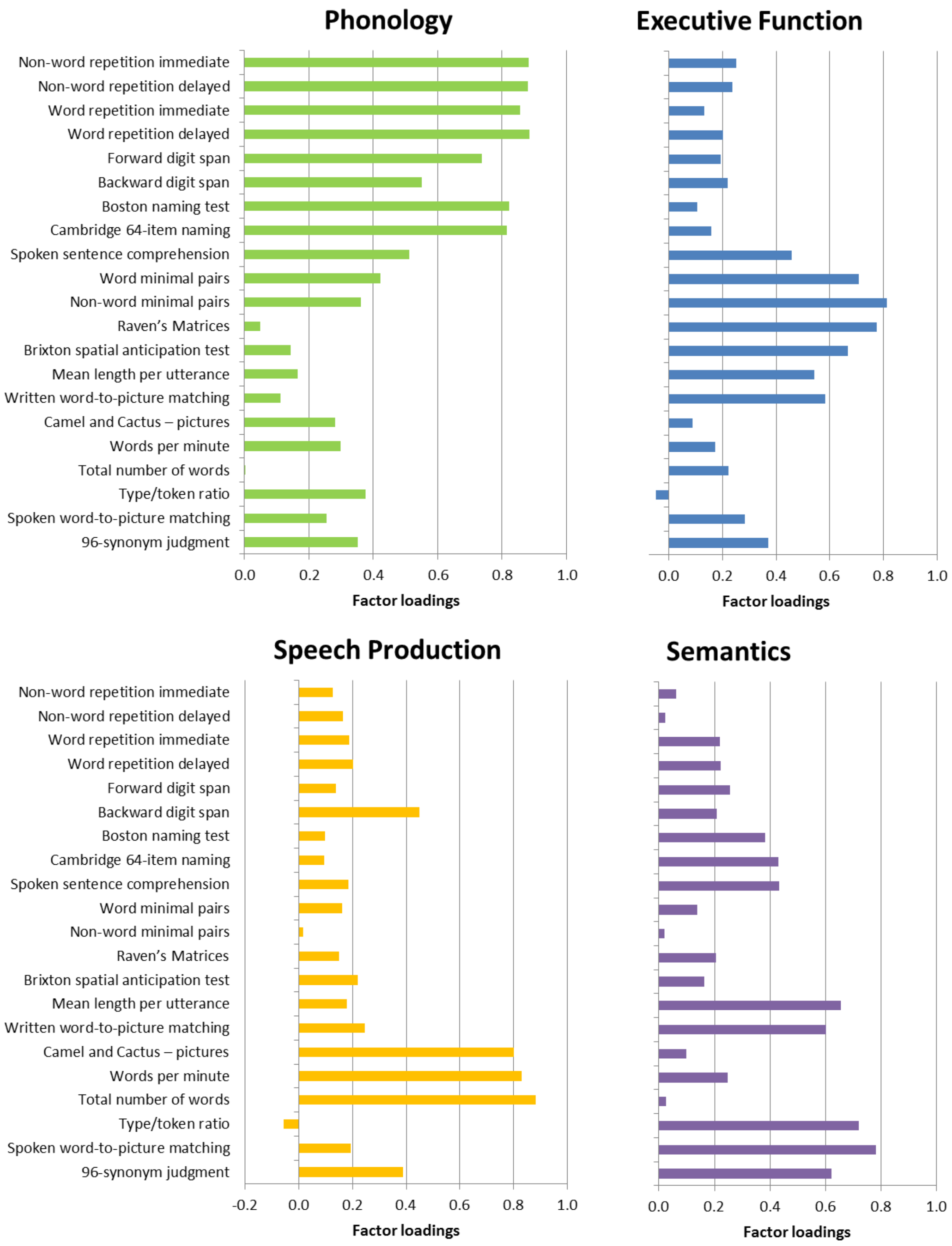


**Figure 1 - Principal components extracted for post-stroke aphasia. Connected speech was elicited using the 'cookie theft' picture description task from the BDAE (Kaplan, 1983). Connected speech was quantified in the parameters: total number of words/tokens, type-token ratio, mean length per utterance, and words-per-minute. Comprehension of grammar was assessed with the spoken sentence comprehension task from the Comprehensive Aphasia Test (Swinburn et al., 2004). Subtests from the Psycholinguistic Assessments of Language Processing in Aphasia (PALPA) battery (Kay et al., 1992) were used to assess auditory discrimination (words and non-words minimal pairs), and repetition of words**

and non-words (immediately and after a delay). Semantic abilities were tested with the Cambridge Semantic Battery (CSB) (Bozeat et al., 2000), including the spoken and written version of the word-to-picture matching task, the 64 item naming test, and the Camel and Cactus Test (picture version). Additionally, the Boston Naming Test (BNT) (Goodglass et al., 1983) and the written 96-trial Synonym Judgement test (Jefferies et al., 2009) were included as these are more sensitive to mild semantic deficits. Non-language measures of attention and executive function included digit span (forward and backwards) (Wechsler, 1981), the Brixton Spatial Rule Anticipation Task (Burgess and Shallice, 1997), and Raven's Coloured Progressive Matrices (Raven and Court, 1962).

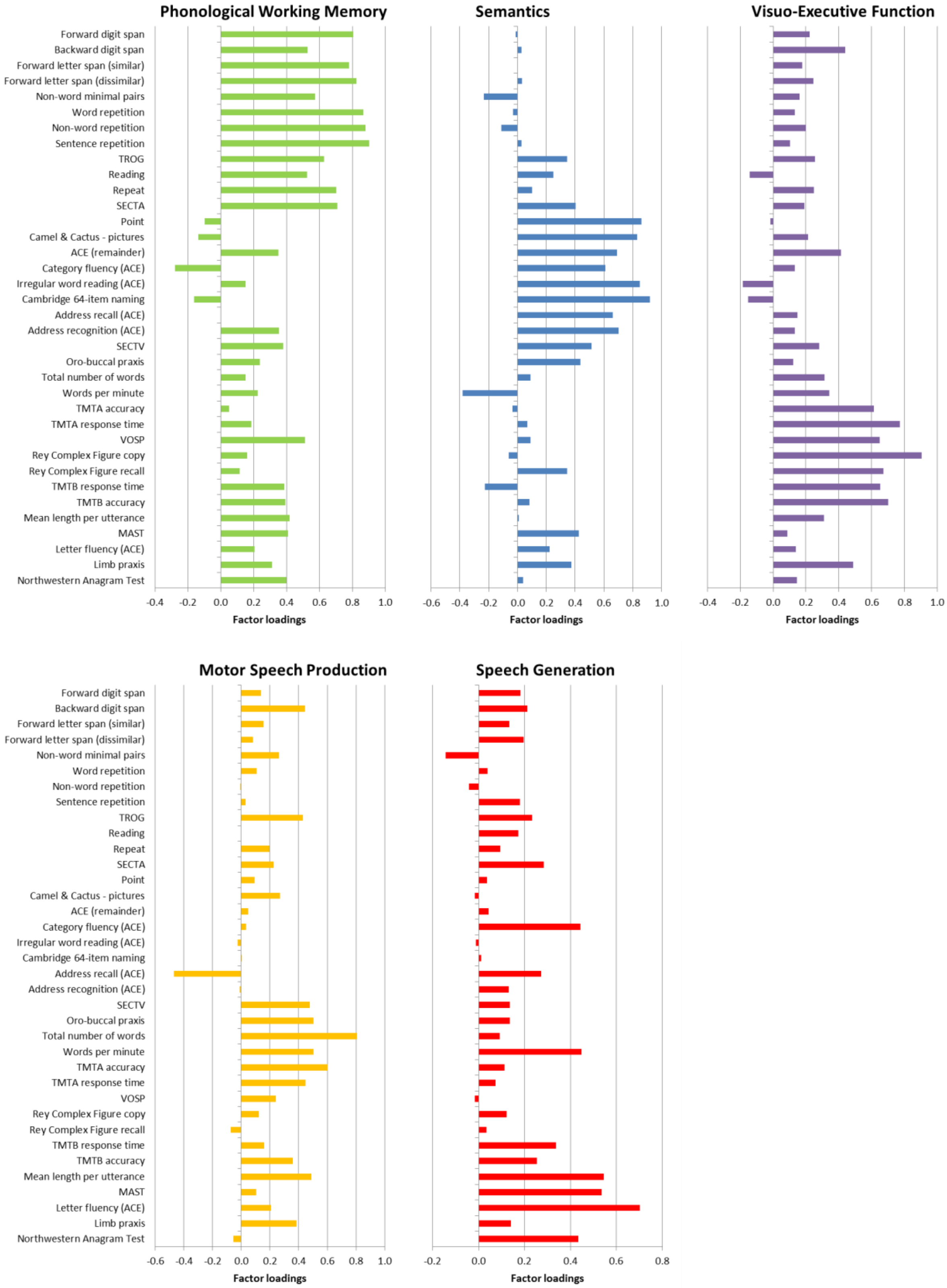


**Figure 2 - Principal components extracted for primary progressive aphasia. Connected speech samples were elicited via picture description task from the Comprehensive Aphasia Test (Swinburn et al., 2004) and quantified in the parameters: total number of words, mean length of utterance (MLU), and words-per-minute (WPM). Comprehension of grammar was assessed through the Test for Reception of Grammar (TROG) (Bishop, 1989) and Sentence Comprehension Test (auditory and visual presentations – SECTA/SECTV) (Billette et al., 2015). The Make A Sentence Test (MAST) (Billette et al., 2015) was used to measure grammatical ability in sentence production. Ability to repeat was tested for words, non-**

words and sentences (Sajjadi et al., 2012). Repetition of words was also measured in the Repeat and Point test (Hodges et al., 2008). Semantic ability was measured through the 64-item naming test and the picture version of the Camel and Cactus Test of semantic association knowledge (CCT), both from the Cambridge Semantic Battery (CSB) (Bozeat et al., 2000). The Point aspect of the Repeat and Point test (Hodges et al., 2008) also measured object knowledge. Phonological skill was assessed through non-word minimal pairs from the Psycholinguistic Assessments of Language Processing in Aphasia (PALPA) (Kay et al., 1992) for phonological perception, and forward letter span (Sajjadi et al., 2012) for measuring the capacity of the phonological loop. Addenbrooke's Cognitive Examination – Revised (ACE-R) (Mioshi et al., 2006) was included as a general assessment which tapped multiple domains of cognition and language. Digit span (Wechsler, 1981) (forwards and backwards) was used to measure attention and executive function, alongside subsections A and B of the Delis-Kaplan executive function system (D-KEFS) Trail Making Test (Delis et al., 2001). Visuospatial skills were tested using the cube analysis subsection of the Visual Object and Space Perception (VOSP) battery (Warrington and James, 1991). The Rey-Osterrieth Complex Figure (Rey, 1941) was also used to measure visuospatial ability in the direct copying of the figure, and non-verbal memory in the later recall of the figure at a delay. Sajjadi *et al.* (2012) also developed tests for oro-buccal (mouth/cheek) and limb praxis, including transitive (meaningful) and intransitive (meaningless) gestures.

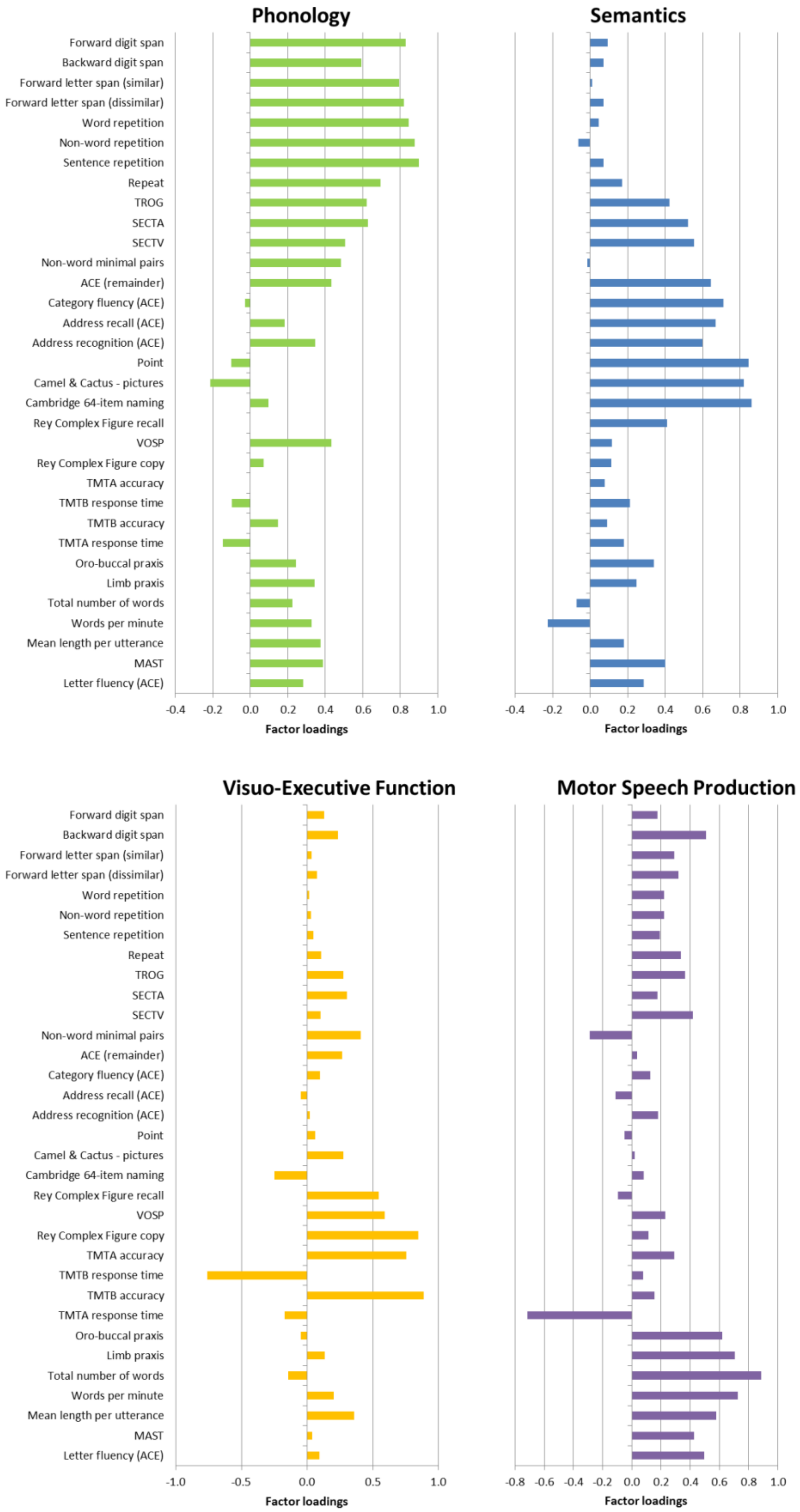


**Figure 3 - Principal components extracted for the unified principal component analysis of primary progressive aphasia and post-stroke aphasia.**

**Table 1 – Results of the analysis to search for categorical subgroups: diagnostic cut-off values derived from iterative sweep through the Unified principal component analysis multi-dimensional space, d prime (d') values associated with each combination of cut-off values, and p-value associated with each d' derived from Monte-Carlo procedure.**

| | | Optimum cut-off values | | | | | |
|---|---|---|---|---|---|---|---|
| Group | Subtype | Phonology | Semantics | Visuo-executive | Speech production | d prime | P value |
| PSA | Anomia | > -1.14 | > 0.38 | > -1.49 | < 1.10 | 2.64 | 0.0001 |
| | Broca | < -0.02 | > 0.33 | > 0.92 | < -0.98 | 2.37 | 0.0012 |
| | Conduction | < -1.22 | > -0.25 | > 0.47 | > -0.88 | 3.11 | 0.0001 |
| | Global | < -1.62 | < -0.88 | > 0.38 | < 0.22 | 3.40 | 0.0001 |
| | Mixed non-fluent | < 1.23 | < 0.39 | > 0.53 | < -1.76 | 2.37 | 0.0019 |
| | TMA | > 0.68 | > 0.54 | > 0.53 | < -1.70 | 2.17 | 0.0288 |
| PPA | lvPPA | > -0.19 | > 0.28 | < -0.59 | > 0.49 | 3.38 | 0.0001 |
| | mPPA | < 1.51 | < 0.73 | < 0.54 | < 1.35 | 2.10 | 0.0001 |
| | PNFA | < 1.37 | > 0.07 | < 0.73 | < 1.83 | 2.03 | 0.0001 |
| | SD | > 0.29 | < 0.98 | > -0.14 | > -0.52 | 4.46 | 0.0001 |

**Table 2 - Distribution of misclassifications between clinical and data-driven diagnostic PSA groups. The cut-off values giving optimum sensitivity for each diagnostic group were treated as data-driven diagnostic criteria. Rows represent 'real' clinical diagnostic categories. The 'Hits' column represents the percentage of patients meeting the data-driven cut-off values for their own data-driven diagnostic group. The columns under 'Misclassifications' represent the percentage of cases whose factor scores (a) met the cut-off values for a different data-driven diagnostic group; (b) did not meet the cut-off values for any of the data-driven diagnostic groups; (c) met the cut-off values for more than one data-driven diagnostic group. These 'Misclassifications' columns are not mutually exclusive, so row totals do not add up to 100%.**

| | | Data-driven diagnostic groups | | | | | | | | |
|---|---|---|---|---|---|---|---|---|---|---|
| | | Hits | Misclassifications | | | | | | | |
| | | | Anomia | Broca | Conduction | Global | Mixed non-fluent | TMA | None | >1 |
| Clinical diagnostic groups (N) | Anomia (15) | 100.0 | / | 0.0 | 0.0 | 0.0 | 0.0 | 0.0 | 0.0 | 0.0 |
| | Broca (5) | 60.0 | 60.0 | / | 0.0 | 0.0 | 0.0 | 0.0 | 20.0 | 40.0 |
| | Conduction (3) | 100.0 | 0.0 | 0.0 | / | 0.0 | 0.0 | 0.0 | 0.0 | 0.0 |
| | Global (5) | 100.0 | 0.0 | 0.0 | 0.0 | / | 0.0 | 0.0 | 0.0 | 0.0 |
| | Mixed non-fluent (5) | 60.0 | 0.0 | 0.0 | 0.0 | 0.0 | / | 0.0 | 40.0 | 0.0 |
| | TMA (1) | 100.0 | 100.0 | 0.0 | 0.0 | 0.0 | 0.0 | / | 0.0 | 100.0 |

Amongst the PSA cases, data-driven diagnostic criteria for three subtypes achieved a perfect hit rate and no misclassifications: global aphasia, conduction aphasia, and TMA subtypes. The data-driven diagnostic criteria for global aphasia and conduction aphasia had the highest d' values (global: d' = 3.40, $p < .001$; conduction: d' = 3.11, $p < .001$), whilst the TMA criteria had a lower d' value (d' = 2.17, p = 0.029). These d' values are lower than that for SD due to the smaller sample sizes in the PSA cohort, which affect the adapted d' calculation (Macmillan and Kaplan, 1985). The anomia data-driven diagnostic group had a perfect hit rate but misclassifications of Broca's aphasia and TMA cases resulting in a lower d' value (d' = 2.64, $p < .001$). The data-driven diagnostic criteria for Broca's aphasia incurred no misclassifications of other subtypes but failed to correctly classify all the Broca's cases (Hit rates around 60%), resulting in lower d' values (Broca: d' = 2.37, p = .001). This shows that, like PNFA, even the optimal data-driven diagnostic criteria for Broca's aphasia and were insufficiently selective, suggesting that this subtype of PSA does not meet the assumptions of a true category.

The mixed non-fluent aphasia group represents cases who did not meet the criteria for a single proposed subtype of PSA. The data-driven diagnostic criteria for this group (d' = 2.37, p = .002) were poorly selective, as they failed to capture all the mixed non-fluent PSA cases. This reinforces the label applied to these cases.
